# A unified population-genetic framework for inferring double reduction and higher-order two-locus linkage disequilibrium in autopolyploids

**DOI:** 10.64898/2026.05.01.722183

**Authors:** Weihao Dou, Zhongfan Lv, Dongchao Ji, Chonghui Zhao, Na Wang, Feng tang Yang, Libo Jiang

## Abstract

Polyploid genomes exhibit complex allelic interactions driven by polysomic inheritance that cannot be adequately captured by conventional pairwise linkage disequilibrium (LD) models, leaving higher-order dependency structures largely unresolved. Here, we develop a unified population-genetic framework that jointly infers double reduction and higher-order allelic associations in autopolyploids. By modeling dosage-dependent gametic structure, our approach reformulates LD as a hierarchical system, enabling decomposition of allelic dependencies beyond pairwise interactions. Through simulations, we show that conventional estimators systematically confound higher-order allelic structure with pairwise LD, leading to biased inference and loss of identifiability under dosage uncertainty. In contrast, our joint likelihood framework achieves asymptotically unbiased estimation and reveals that effective double reduction emerges as a composite parameter linking meiotic configuration to population-level allelic reshaping. Applying this model to *Arabidopsis arenosa* and cultivated potato, we uncover fundamental contrasts in genomic architecture. In natural populations, higher-order LD self-organizes into structured, long-range dependency networks, whereas predominantly clonal populations exhibit collapse of higher-order dependency structure. We further identify a spatial antagonism between double reduction and higher-order LD, demonstrating that elevated double reduction compresses the physical scale of allelic interactions. Our results demonstrate that autopolyploid genomes are organized as higher-order dependency systems rather than collections of pairwise associations, establishing higher-order allelic structure as a fundamental extension of classical LD theory.

## Introduction

Polyploidization is a major driver of plant evolution, shaping speciation, environmental adaptation, and the domestication of many important crops (Van de Peer *et al*, 2021; Heslop-Harrison *et al*., 2023). Numerous economically and ecologically significant species are polyploid (Rice *et al*., 2019; Fox *et al*., 2020). Polyploids are broadly classified into two major categories, autopolyploids and allopolyploids, which differ fundamentally in their genomic composition and meiotic behavior (Bever & Felber, 1992). In autopolyploids, multivalent pairing and recombination among multiple homologous chromosomes during meiosis lead to polysomic inheritance and highly complex segregation patterns (Grandont *et al*., 2013; Lv *et al*., 2024). These processes generate intricate patterns of allelic association across the genome, yet current population-genetic frameworks largely characterize linkage disequilibrium (LD) in terms of pairwise relationships between loci (Jiang *et al*., 2021; Gerard, 2021). Whether polyploid genomes are instead organized by higher-order allelic dependencies, arising from polysomic inheritance and recombination, remains largely unexplored.

A key cytological feature of autopolyploids is double reduction (DR), in which sister chromatid segments from the same homolog co-segregate into a single gamete (Darlington, 1929). DR has long been recognized to alter genotype frequencies at individual loci, and recent studies have developed statistical frameworks to estimate its occurrence in natural autotetraploid populations, assuming Hardy-Weinberg equilibrium (HWE) (Jiang *et al*., 2021). However, how the effects of DR propagate from single-locus genotype frequencies to multi-locus allelic associations remains unclear. In particular, it remains unknown whether DR systematically reshapes the structure of linkage disequilibrium beyond pairwise interactions, potentially generating higher-order dependency patterns that cannot be captured by existing models.

Linkage disequilibrium quantifies the non-random association of alleles across distinct loci within a natural population. As a sensitive indicator of population genetic forces, the extent and genomic distribution of LD provide key insights into demographic history, evolutionary processes, and the genetic dissection of complex traits (Wu & Zeng, 2001; Sved & Hill, 2018; Harris *et al*., 2025). While LD models developed for diploids, such as haplotype-based approaches, have been widely applied in population-genetic analyses, their underlying assumptions limit these model’s ability to capture the complex structure of LD in polyploid gametes (Dufresne *et al*., 2014; Meirmans *et al*., 2018). This limitation reflects the nature of polysomic inheritance in autopolyploids, which generates multi-allelic gametes, dosage-dependent interactions among homologous chromosomes, and a high-dimensional genotype space (Huang *et al*., 2022). As a result, polyploid genomes may exhibit higher-order allelic dependencies that cannot be reduced to pairwise LD. However, most current methods rely on pairwise LD statistics (Gerard, 2021), which fail to resolve these higher-order structures. Moreover, existing approaches do not explicitly incorporate double reduction (DR) or Hardy-Weinberg disequilibrium (HWD), instead either assuming random bivalent pairing or subsuming their effects into deviations from equilibrium (Gerard, 2023). Whether such confounding extends beyond single loci to genome-wide allelic dependency structures remains unclear.

Zygotic disequilibria, originally developed for diploid populations (Cockerham and Weir, 1977), have been extended to polyploids through diploid-based representations, but do not generalize to autopolyploids (Yang *et al*., 2022), where polysomic inheritance and DR reshape allelic association patterns. Here, we developed a population-genetic framework to quantify higher-order allelic dependencies in autopolyploids. The framework captures dosage-dependent interactions among homologous chromosomes and accommodates different ploidy levels, including autotetraploid (4x) and autohexaploid (6x) systems. It enables parameter inference under incomplete genotype dosage information and is implemented in an open-source package, **AutoLD**. We applied this approach to chromosome-scale data from species representing distinct population structures, including cultivated potato (*Solanum tuberosum*) and natural populations of *Arabidopsis arenosa*. By separating the effects of meiotic processes from population-level associations, the framework provides a general basis for analyzing genetic structure and evolutionary dynamics in autopolyploids. In particular, we introduce the concept of an effective double reduction parameter (*α*_*eff*_), which integrates meiotic and population-level processes into a unified statistical quantity.

## Result

### Joint inference remains robust under dosage uncertainty and model misspecification

Joint inference of higher-order allelic dependencies remains statistically identifiable under both equilibrium and disequilibrium conditions. We evaluated the statistical properties of our joint inference framework through extensive simulations under both Hardy-Weinberg equilibrium (HWE) and disequilibrium (HWD). Simulation designs are summarized in Supplementary Tables S1-S2. Across experiments, we assessed empirical false positive rates (FPR), statistical power, and estimation bias as functions of sample size and marker informativeness, comparing dosage-resolved (R) with dosage-unresolved (U) genotypes.

Under the null (Linkage equilibrium, HWE; Simulation series I; Table S3), the framework maintained nominal Type I error when full dosage information was available. In autotetraploids at *N* = 500, empirical FPRs for both double reduction coefficients and LD parameters clustered around 0.05. For instance, pairwise LD (*D*_*eab*_) showed FPR = 0.05, while higher-order terms yielded 0.06 and 0.07, respectively. Autohexaploids exhibited similarly well-behaved FPRs (0.00-0.06). Dosage ambiguity, however, induced marked conservatism. In autotetraploids at *N* = 500, FPR for *D*_*eab*_ dropped to 0.01, and most of the higher-order interactions approached zero. This collapse was more severe in autohexaploids, where essentially all LD parameters yielded FPR = 0.00 across sample sizes (Table S3). Loss of dosage resolution substantially reduces the information available for LD inference, producing flatter likelihood surfaces and increasingly conservative test behavior.

We next evaluated whether strong pairwise LD could inflate higher-order interaction signals (Linkage equilibrium, HWE; Simulation series II; Table S4). We fixed background pairwise linkage at *D*_*eab*_ = 0.05 while setting all higher-order terms to zero. Under HWE, FPRs for higher-order parameters remained tightly controlled. In autotetraploids at *N* = 500, *D*_*Ab*_, *D*_*aB*_, and *D*_*AB*_ yielded FPRs of 0.05, 0.06, and 0.06, respectively. Autohexaploids showed even lower FPRs-typically <0.05 across all parameters, including the sixth-order term *D*_*AABB*_ (FPR = 0.05 at *N* = 1000). The decomposition successfully isolates hierarchical association structure: higher-order components capture variance orthogonal to pairwise LD, not mere aliasing.

Statistical power to detect nonzero LD and the precision of estimates both depended on genotype resolution (Linkage equilibrium, HWE; Simulation series III; Table S5). For autotetraploid pairwise LD (*D*_*eab*_ = 0.050), full dosage information at *N* = 500 yielded an unbiased MLE (0.050 ± 0.012) with power 0.96. Under dosage ambiguity, the estimate remained nearly unbiased (0.053 ± 0.025), but power plummeted to 0.59. This resolution gap widened for higher-order terms. Detecting the third-order interaction *D*_*Ab*_=0.020 at *N* = 500 achieved power 0.97 under resolved data versus 0.62 under unresolved; for the fourth-order term *D*_*AB*_=0.010, power dropped from 0.88 to 0.28 (Table S5). In autohexaploids, higher-order parameters were intrinsically harder to estimate. Even at *N* = 1000, power for the sixth-order interaction *D*_*AABB*_ = 0.001reached only 0.75 with resolved dosage and 0.34 without. Despite the severe loss of statistical power under partial dosage, the maximum likelihood estimates (MLEs) remained asymptotically unbiased across all settings.

Real populations may deviate from the assumption of no double reduction. We therefore assessed robustness by simulating increasing true *α* while fitting the misspecified model *α* = 0 (Linkage equilibrium, HWE; Simulation series IV; Table 1). In autotetraploids, pairwise LD estimation remained highly stable even when the true double reduction rate approached its theoretical upper bound. For example, at *N* = 500, the power to detect *D*_*eab*_ remained above 0.95 across the tested range of *α*, and estimation bias remained negligible (Table 1). However, inference in autohexaploids was more sensitive to segregation misspecification. As the true double reduction rate increased, the stability of high-order LD estimation deteriorated, with increasing false positive rates and reduced statistical power for some parameters. The denser association space in hexaploids amplifies the consequences of segregation misspecification, serving as a critical caution against applying bivalent-inspired models to complex autopolyploids.

**Table 1.** Robustness of dosage-resolved LD estimation to model misspecification under Hardy-Weinberg equilibrium across ploidy levels.

| | $\alpha = 0.000$ | | | $\alpha = 0.050$ | | | $\alpha = 0.100$ | | | $\alpha = 0.167$ | | |
| --- | --- | --- | --- | --- | --- | --- | --- | --- | --- | --- | --- | --- |
| | FPR | Bias<br>($\times 10^{-3}$) | Power | FPR | Bias<br>($\times 10^{-3}$) | Power | FPR | Bias<br>($\times 10^{-3}$) | Power | FPR | Bias<br>($\times 10^{-3}$) | Power |
| Autotetraploid ($N = 500$) | | | | | | | | | | | | |
| $D_{eab}$ | 0.06 | 0.66 | 0.98 | 0.06 | 2.74 | 0.97 | 0.06 | -6.43 | 0.95 | 0.05 | -10.73 | 0.96 |
| $D_{Ab}$ | 0.05 | 1.74 | 0.94 | 0.07 | 0.88 | 0.93 | 0.07 | -1.25 | 0.93 | 0.05 | -1.51 | 0.90 |
| $D_{aB}$ | 0.06 | 1.51 | 0.58 | 0.06 | 0.13 | 0.49 | 0.05 | -1.54 | 0.36 | 0.05 | -1.77 | 0.31 |
| $D_{AB}$ | 0.06 | 0.27 | 0.79 | 0.07 | 0.26 | 0.69 | 0.04 | 0.57 | 0.71 | 0.05 | 0.53 | 0.71 |
| Autohexaploid ($N = 1000$) | | | | | | | | | | | | |
| $D_{eab}$ | 0.12 | 14.81 | 0.93 | 0.18 | 16.92 | 0.92 | 0.20 | 15.34 | 0.92 | 0.23 | 18.45 | 0.87 |
| $D_{Ab}$ | 0.15 | 4.08 | 0.45 | 0.27 | -5.34 | 0.40 | 0.30 | -6.03 | 0.41 | 0.31 | -6.84 | 0.36 |
| $D_{aB}$ | 0.14 | 5.97 | 0.87 | 0.25 | 6.42 | 0.85 | 0.31 | 6.96 | 0.75 | 0.32 | 7.02 | 0.66 |
| $D_{AAb}$ | 0.13 | 5.94 | 0.76 | 0.20 | 6.31 | 0.70 | 0.33 | 6.75 | 0.60 | 0.37 | 7.05 | 0.48 |
| $D_{aBB}$ | 0.12 | 3.89 | 0.51 | 0.21 | 4.51 | 0.44 | 0.32 | 4.45 | 0.42 | 0.36 | 4.79 | 0.38 |
| $D_{AB}$ | 0.19 | -3.78 | 0.49 | 0.32 | -4.9 | 0.45 | 0.38 | -4.82 | 0.43 | 0.38 | -4.95 | 0.41 |
| $D_{AAB}$ | 0.20 | 2.62 | 0.81 | 0.31 | 3.02 | 0.76 | 0.35 | 3.46 | 0.70 | 0.41 | 4.52 | 0.56 |
| $D_{ABB}$ | 0.19 | -1.88 | 0.41 | 0.32 | -2.77 | 0.42 | 0.40 | -3.22 | 0.37 | 0.40 | -3.95 | 0.39 |
| $D_{AABB}$ | 0.18 | 0.76 | 0.59 | 0.31 | 1.00 | 0.56 | 0.41 | 1.05 | 0.49 | 0.44 | 1.06 | 0.43 |

Note: Data are derived from 1,000 simulations for FPR assessment and autotetraploid analysis, and 500 simulations for autohexaploid Bias/Power analysis. Sample sizes were set to $N = 500$ for autotetraploids and $N = 1000$ for autohexaploids. The estimation model strictly assumes no double reduction ( $\alpha = 0$ ). FPR: False Positive Rate evaluated under the null hypothesis (all $D$ parameters set to 0) at a nominal significance level of 0.05. Bias/Power: Evaluated under alternative hypotheses with fixed linkage parameters. For autotetraploids, true values were $D_{eab} = 0.05$ , $D_{Ab} = 0.02$ , and $D_{aB} = D_{AB} = 0.01$ . For 6x, true values were $D_{eab} = 0.03$ , with higher-order interactions ( $D_{Ab}$ through $D_{AABB}$ ) ranging from 0.001 to 0.005. Bias values are scaled by $10^{-3}$ for clarity. $A = 0.167$ represents the theoretical maximum double reduction rate for autotetraploids (1/6).

Finally, we examined the impact of HWD on parameter identifiability (Table S2). When genotype frequencies deviate from HWE, the estimated double reduction parameter no longer reflects the true meiotic double reduction rate, but instead represents an effective segregation parameter (*α*_*eff*_) that absorbs all sources of disequilibrium, including true double reduction, non-random mating, and population structure. In autotetraploids at *N* = 200, the empirical FPRs for *α*_*effA*_ and *α*_*effB*_ reached 62.9% and 64.3%, respectively, increasing to nearly 100% at *N* = 1000 (Table S6). With sufficient sample size, any HWD-induced distortion is systematically attributed to *α*_*eff*_, rendering true double reduction statistically indistinguishable from other forces that perturb genotype frequencies. Under HWD, FPRs for LD parameters stayed near 0.05 (Table S6). Although statistical power was moderately reduced, dropping from 0.96 to 0.82 for *D*_*eab*_ at *N* = 500, the MLEs remained unbiased (Table S7). This asymmetry arises from the hierarchical structure of the likelihood, where allelic association components occupy a subspace orthogonal to the deviations absorbed by *α*_*eff*_. Because of this structural decoupling, severe HWD precludes reliable inference about meiotic mechanisms like double reduction, yet leaves the detection of linkage-driven associations entirely uncompromised.

### Higher-order gametic dependence biases conventional LD estimators

Higher-order gametic dependence systematically biases conventional LD estimators when unmodeled. We compared the proposed joint inference framework (AutoLD) against three alternative estimators, including Dip (diploid approximation), Hap (haplotype-based), and ldsep, under simulations that progressively introduced higher-order biological complexity while holding pairwise LD constant (*D*_*eab*_ = 0.05; Figure 1A). Three scenarios were considered: pairwise LD only (S1), pairwise plus higher-order LD (S2: HOLD), and a full polysomic model with both higher-order LD and locus-specific double reduction (S3: HOLD + DR).

**Figure 1.**
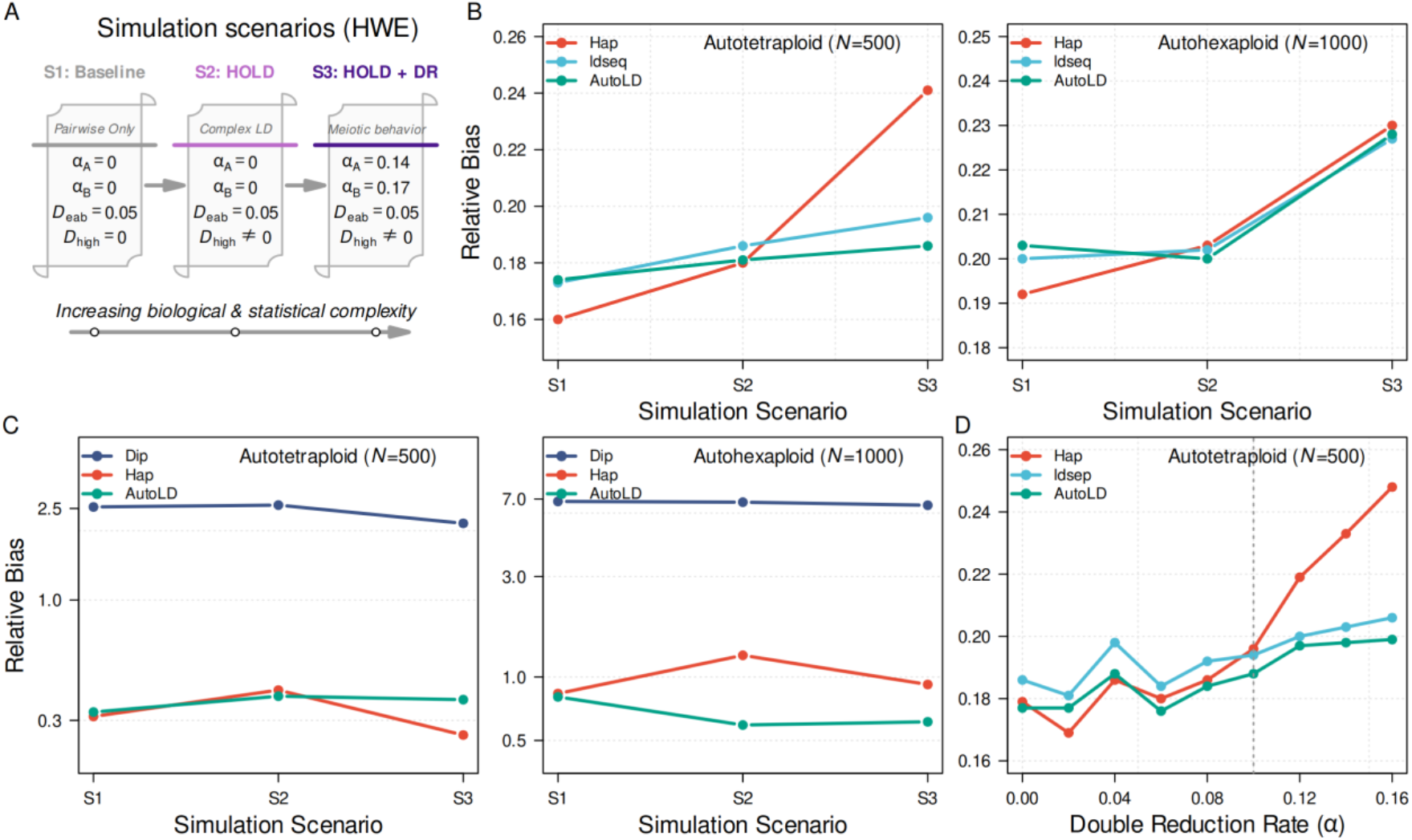
Benchmarking LD estimation models under simulated polyploid scenarios. (A) Simulated populations under Hardy-Weinberg equilibrium with increasing genetic complexity. Three scenarios are considered: absence of higher-order LD and double reduction (S1); structured multiallelic dependence (S2: HOLD); and combined higher-order LD and double reduction (S3: HOLD+DR). (B-C) Accuracy of LD estimation across autotetraploid (*N* = 500) and autohexaploid (*N* = 1000) genomes, evaluated by relative bias. Methods include the diploid approximation (Dip), haplotype model (Hap), a polyploid LD estimator (ldsep), and the proposed method (AutoLD). (D) Relative bias of LD estimates as a function of double reduction rate (*α*) in autotetraploids (*N* = 500).

When only pairwise LD was present (S1), all dosage-aware estimators performed similarly. In autotetraploids (*N* = 500), relative bias for Hap, ldsep, and AutoLD ranged from 0.16 to 0.18; in autohexaploids (*N* = 1000), all three clustered around 0.20 (Figure 1B). The addition of higher-order LD (S2) revealed divergence. Hap bias increased to 0.18-0.19 in autotetraploids, ldsep to 0.19-0.20, while AutoLD remained stable near 0.18. This gap widened under the full meiotic model (S3): in tetraploids, Hap bias reached ∼0.24, ldsep ∼0.20, whereas AutoLD stayed below 0.19. Hexaploids showed a similar pattern under S3, Hap bias climbed to ∼0.23, AutoLD held near 0.22 (Figure 1B). Conventional estimators increasingly misattribute higher-order gametic dependence to pairwise LD.

When genotype dosage was partially collapsed (Figure 1C), the diploid approximation (Dip) became severely biased-relative bias exceeded 2.5 in tetraploids and approached 7.0 in hexaploids across all scenarios. Even haplotype-based inference (Hap), though polyploid-aware, showed systematic inflation under dosage uncertainty. AutoLD, in contrast, maintained stable estimates, with relative bias below 0.35 in tetraploids and below 0.60 in hexaploids (Figure 1C). Dosage resolution is not merely a technical nuance; it fundamentally determines whether higher-order dependence can be distinguished from pairwise signal.

We further examined estimator behavior across increasing double reduction rates while maintaining higher-order LD (Figure 1D). As the double reduction parameter increased from *α* = 0 to approximately 0.16, the relative bias of conventional estimators increased steadily. In autotetraploids, the relative bias of Hap rose from ∼0.17 to ∼0.25. The composite estimator ldsep showed a similar upward trend. In contrast, the bias of AutoLD remained nearly constant across the entire range of double reduction values, staying close to 0.20 (Figure 2D). This stability indicates that explicitly modeling segregation parameters prevents double reduction from being absorbed into the pairwise LD estimate.

**Figure 2.**
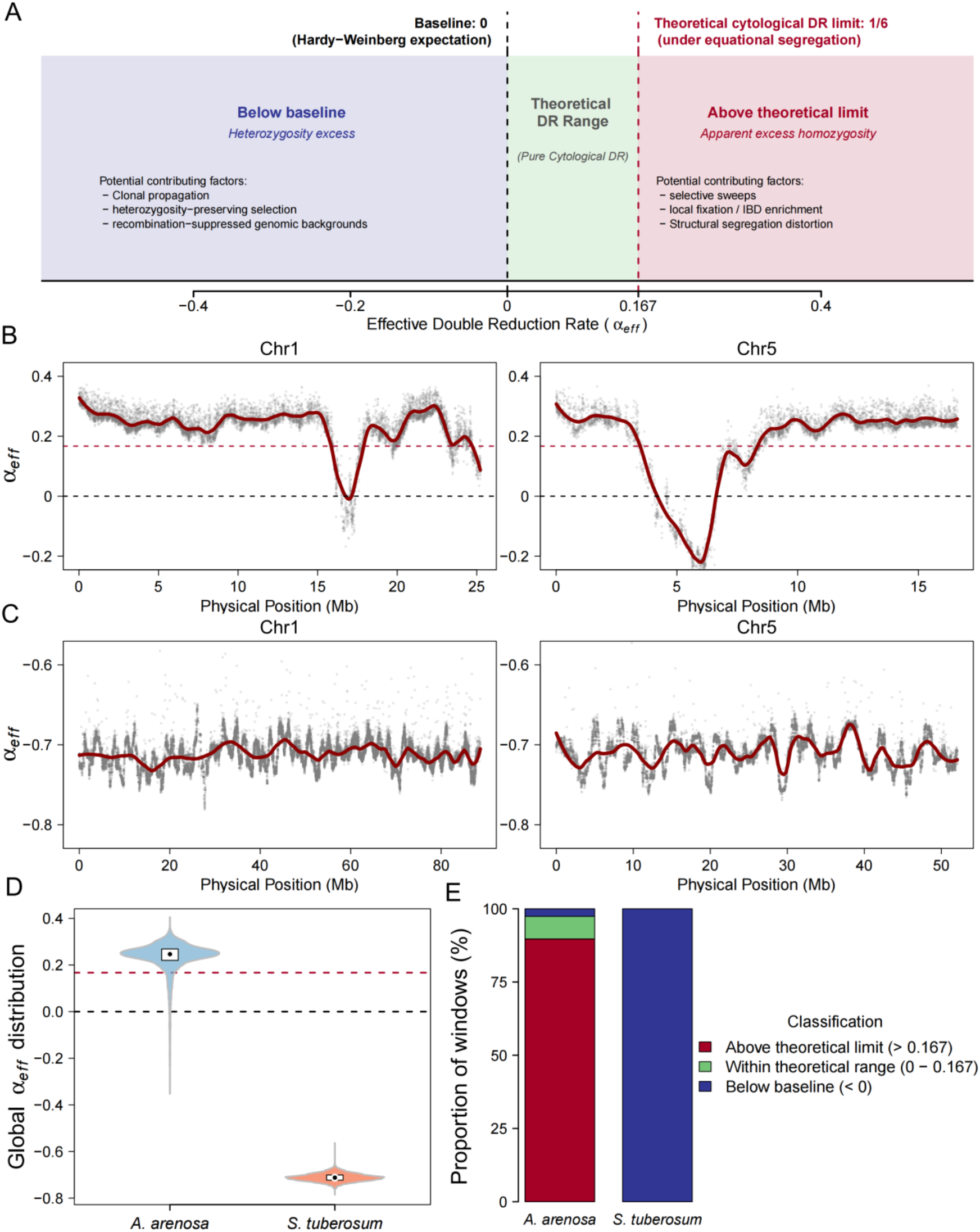
The chromosome-scale variation of effective double reduction (*α*_*eff*_) across autopolyploid populations. (A) Conceptual framework for estimating *α*_*eff*_ in autotetraploids. Three regions are defined: expected meiotic range (0 ≤ *α*_*eff*_ ≤ 1/6), excess homozygosity (*α*_*eff*_ > 1/6), and excess heterozygosity (*α*_*eff*_ < 0). (B-C) Spatial variation of *α*_*eff*_ along representative chromosomes (Chr1 and Chr5) for *A. arenosa* (B) and *S. tuberosum* (C). The horizontal black dashed line indicates *α*_*eff*_ = 0, the red dashed line denotes the genome-wide mean in *A. arenosa*. LOESS curves are shown in dark red. (D-E) Distribution of *α*_*eff*_ across the genome (D) and the proportion of genomic windows across the three theoretical regions (E) in *A. arenosa*.

Because HWD produces genotype frequency distortions that mimic double reduction, we repeated the simulations under HWD (Figure S1). Bias inflation for conventional estimators became more pronounced. In hexaploids, Hap bias increased from ∼0.36 in S1 to ∼0.43 in S3, while AutoLD remained stable around 0.27 (Figures S1B). The same pattern held across increasing *α* (Figure S2D). HWD does not create the bias, but it amplifies the vulnerabilities of estimators that fail to account for higher-order gametic structure. These results demonstrate that while segregation distortions systematically confound standard LD metrics, the joint likelihood formulation of AutoLD successfully isolates these effects, maintaining a stable and unbiased estimation of the structural two-locus LD component.

### Effective double reduction captures meiotic and population-level signals in natural and cultivated autotetraploid populations

Effective double reduction integrates meiotic and population-level processes into a unified genomic signal. To clarify the meaning of the combined parameter *α*_*eff*_, we first created an analytical framework. This framework was designed to explain how *α*_*eff*_ is expected to behave under different genetic conditions (Figure 2A). In this scheme, *α*_*eff*_ = 0 corresponds to the Hardy-Weinberg baseline under random mating, while the theoretical upper bound for meiotic double reduction in autotetraploids under complete equational segregation is 1/6. Estimates falling within this interval are thus compatible with the cytologically permissible range of double reduction; values outside this range signal the influence of additional population-genetic forces shaping genotype frequencies.

Guided by this interpretive scaffold, we applied the model to two autotetraploid populations chosen for their sharply contrasting evolutionary and reproductive backgrounds: a naturally outcrossing population of *Arabidopsis arenosa* and a highly domesticated lineage of *Solanum tuberosum* (cultivated potato). The former represents a wild, sexually propagating polyploid system (Monnahan *et al*., 2019); the latter has undergone intensive artificial selection coupled with long-term clonal propagation-a dichotomy that provides a powerful context for interpreting the multifaceted signals captured by *α*_*eff*_. For the potato analysis, we drew on six chromosome-scale assemblies currently available in the NGDC database, whose contiguous structure enables high-resolution tracking of the statistic along individual chromosomes.

In *A. arenosa*, the genomic profile of *α*_*eff*_ provided a comprehensive view of meiotic limitations. Along the distal arms of chromosomes, smoothed trajectories consistently exhibited elevated levels, with local fluctuations focused around positive values (Figure S2). Conversely, these trajectories sharply declined in areas corresponding to inferred pericentromeric intervals, resulting in typical V- or U-shaped patterns (Figure 2B). This spatial gradient corresponds with the established restriction of recombination at centromeres, which physically restricts the potential for twofold reduction. Consequently, at the chromosomal level, αev accurately reflects traditional cytological predictions, mapping the gradient in crossover frequency from centromere to telomere.This spatial gradient corresponds to the established suppression of recombination near centromeres, which physically restricts the potential for double reduction (Mather, 1935; Bourke *et al*., 2017).Thus, at the chromosomal level, αev effectively mirrors established cytological expectations, delineating the gradient of crossover frequency from centromere to telomere.

At the genome-wide level, the distribution of *α*_*eff*_ in *A. arenosa* was strongly shifted toward positive territory (Figure 2D). The median value (∼0.25) not only exceeded the Hardy-Weinberg baseline of zero but also surpassed the theoretical cytological maximum for complete equational segregation (1/6 ≈ 0.167). Nearly 90% of genomic windows fell into the “above theoretical limit” category (Figure 2E). This pervasive excess, however, should not be read as a violation of cytological constraints. Because *α*_*eff*_ integrates double reduction with other departures from random mating, the signal likely captures the combined footprint of demographic history and selection. Previous population-genomic work has documented rapid postglacial range expansion and widespread selective sweeps in *A. arenosa* (Monnahan *et al*., 2019), processes that elevate homozygosity through identity-by-descent enrichment across the genome.

The contrast with cultivated potato was striking. Across all analyzed chromosomes, *α*_*eff*_ was globally displaced toward negative values (Figure 2C; Figure S3). The genome-wide median plummeted to approximately -0.71, a dramatic shift relative to *A. arenosa* (Wilcoxon rank-sum test, P < 0.01; Figure 2D). At the window level, every evaluated region fell into the “below baseline” category (*α*_*eff*_ < 0; Figure 2E). This widespread negative signal indicates a genome-wide surplus of heterozygosity compared to Hardy-Weinberg equilibrium, a characteristic of the potato’s reproductive strategy. The sustained clonal reproduction, combined with significant artificial selection for heterotic traits, has promoted the retention of heterozygous genotypes (Hardigan *et al*., 2017; Zhu *et al*., 2025). Without frequent meiotic recombination, these allelic combinations endure over long periods, thereby causing the substantial negative deviation of *α*_*eff*_. This cross-species contrast highlights the dual interpretability of *α*_*eff*_. At the chromosome scale, the statistic reflects meiotic constraints such as recombination suppression around centromeres. At the population scale, it captures broader evolutionary and breeding processes, including demographic expansion and clonal domestication in cultivated potato.

### Higher-order two-locus allelic topology reveals structured dependency organization in autotetraploid genomes

Conventional two-locus linkage disequilibrium analysis in polyploids rests on second-order covariance between allelic states (Gerard, 2021). But this pairwise framing carries an inherent limitation: it collapses over the dosage configurations that define polyploid genomes. Two genotypic arrangements A-A-b-b and A-a-A-a, for instance, can yield nearly identical pairwise signals while representing fundamentally different biological states. The ambiguity is not merely technical; it reflects a loss of information about how alleles are distributed across homologous copies. We therefore extended the covariance framework to third- and fourth-order terms, constructing a tensor-based representation of two-locus associations (Figure 3A). This model accounts for how dosage affects things differently and clearly shows how multiple versions of a gene are arranged on all matching chromosomes. Unlike traditional methods that only measure how closely related things are, the tensor framework describes the structure of these relationships. This provides a more detailed view of the complex nature of polyploid genomes.

**Figure 3.**
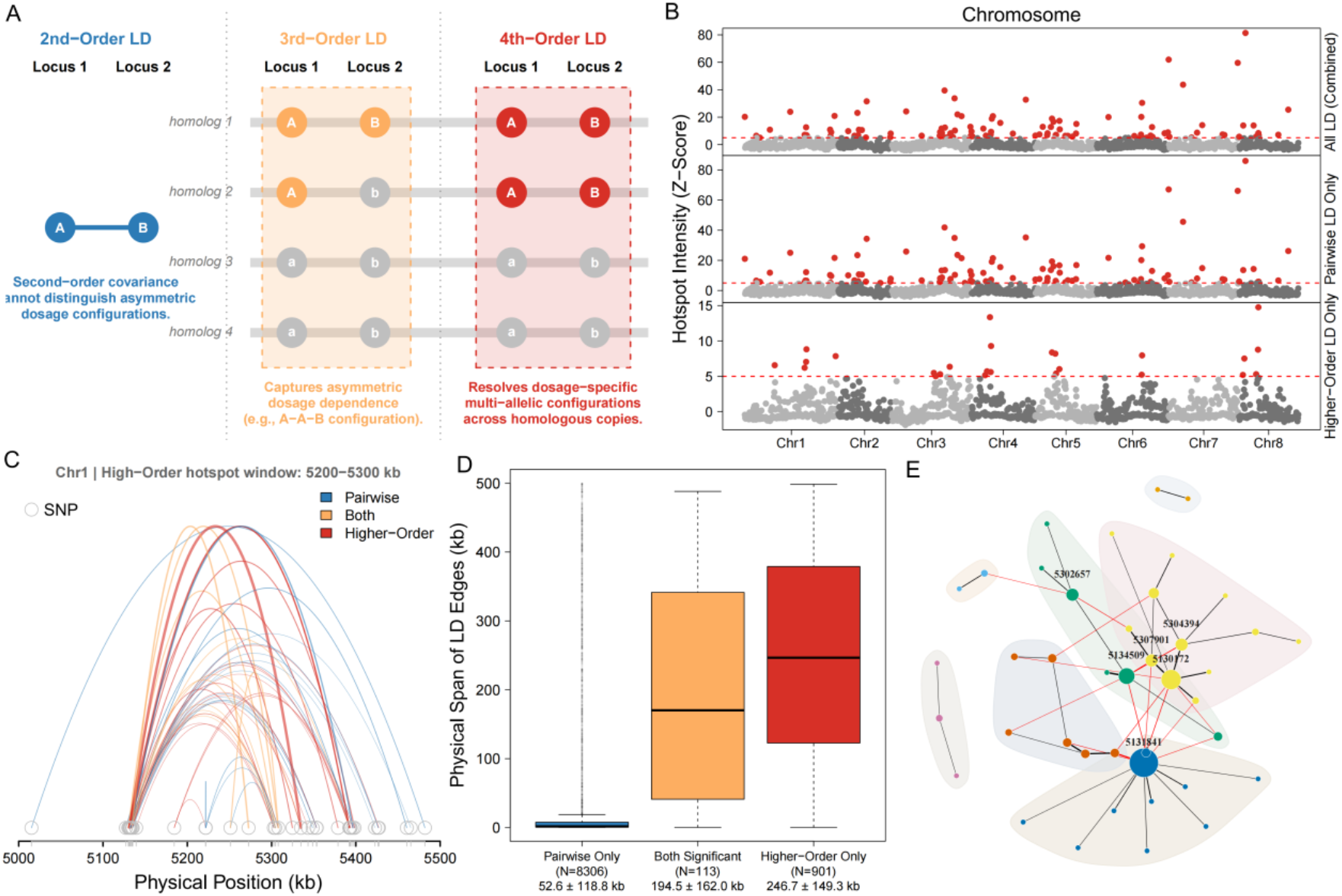
Higher-order topological LD reveals structured networks of long-range allelic dependencies. (A) Comparison of second-order and higher-order LD representations. Second-order covariance cannot distinguish dosage configurations, whereas higher-order LD captures asymmetric allelic combinations across homologous chromosomes. (B) Genome-wide LD hotspot landscape in *A. arenosa*. Red dots represent significant hotspots (*Z* > 5, permutation tests). (C) Local allelic association network within a representative hotspot (Chr1: 5200-5300 kb). Blue edges denote second-order LD; red edges denote higher-order LD. (D) Distribution of LD span lengths across significance levels. (E) Network topology of LD hubs. Node size reflects connectivity; edges indicate significant allelic associations.

Applying this framework to *A. arenosa*, we scanned the genome for regions enriched in higher-order associations (100 kb windows; Z-score ≥ 5). The resulting landscape revealed a sharp divergence between second-order and higher-order LD structures (Figure 3B). Pairwise LD hotspots were broadly distributed, tracing the expected background of short-range linkage. Higher-order hotspots told a different story: they were markedly fewer, spatially confined, and organized into discrete clusters rather than a genome-wide continuum. This spatial decoupling implies that higher-order dependencies do not simply echo pairwise signals. They define specific genomic neighborhoods where allelic configurations are intrinsically more complex.

To examine the internal structure of these regions, we performed a detailed analysis of a representative hotspot on chromosome 1 (5200-5300 kb). Arc-network visualization clearly distinguished between pairwise and higher-order interactions (Figure 3C). Pairwise linkage disequilibrium (LD) edges, depicted as blue arcs, were primarily restricted to short genomic distances, frequently connecting directly adjacent single nucleotide polymorphisms (SNPs). Higher-order edges, represented by red arcs, often encompassed larger physical distances, linking variants that were not identified through simple pairwise associations. These observed patterns imply that higher-order LD incorporates information across multiple loci within the studied region, thereby constructing a network of dependencies that are not discernible from pairwise relationships alone.

To quantify this observation genome-wide, we measured the physical span of LD edges (Figure 3D). Pairwise-exclusive edges averaged 52.6 ± 118.8 kb. Edges significant in both pairwise and higher-order tests spanned 194.5 ± 162.0 kb. Strictly higher-order edges extended further still: 246.7 ± 149.3 kb. The pattern held across independent hotspots (Figure S4), ruling out local artifacts. But span is only part of the story. Within hotspots, these long-range connections coalesced into hub-like structures (Figure 3E). Graph-theoretical metrics confirmed the visual impression: compared to matched null regions, higher-order hotspots showed elevated network transitivity (*P* = 7.6 × 10^-9^) and a pronounced giant component (*P* = 7.8 × 10^-9^) (Figure S5). Higher-order LD in *A. arenosa* is not randomly scattered; it self-organizes into connected modules spanning multiple loci.

Does this architecture depend on how a population reproduces? We turned to cultivated potato (*S. tuberosum*), an autotetraploid propagated clonally for millennia, and applied the same analytical framework. In potato, the genome-wide landscape of higher-order LD was essentially flat (Figure S6A). Pairwise signals remained detectable (Z-score ≥ 5), but the higher-order layer showed sparse, near-absent signals across all chromosomes. Arc-network visualization of a matched genomic region (Chr3: 50997-51540 kb) confirmed the pattern: short-range pairwise edges dominated, with little evidence of higher-order hubs (Figure S6B). The complex dependency modules characteristic of sexually reproducing *A. arenosa* are largely absent under clonal propagation. To ensure this contrast reflected biology rather than metric artifact, we compared multiple LD measures across the same windows (Figure S7). The 95th percentiles told a consistent story: both the higher-order tensor metric and conventional additive dosage-based LD collapsed to similarly low baselines (∼0.01) in potato, while second-order LD retained a modest upper bound (∼0.11). The absence of higher-order signals in potato is not a quirk of the estimator; it is a property of the population. The tensor framework thus provides a generalizable approach for dissecting allelic dependency topology in polyploid genomes, revealing how reproductive mode shapes the architecture of genetic variation.

### Double reduction reshapes LD decay and effective recombination scale in autopolyploid genomes

Double reduction acts as a primary force that reshapes the spatial scale of allelic dependencies in autopolyploid genomes. To evaluate the localized impact of meiotic multivalent pairing on genomic architecture, we exclusively focused on the sexually reproducing *A. arenosa* cohort. Cultivated potato (*Solanum tuberosum*) was deliberately excluded from this specific spatial analysis; its sustained clonal propagation circumvents meiosis entirely, thereby neutralizing double reduction (DR) as an active driver in shaping contemporary linkage disequilibrium (LD) topology.

We first established the genome-wide spatial baseline for allelic dependencies (Figure 4A). Unlike traditional second-order covariance, which exhibits a standard exponential decay initiating near *r*^2^ ≈ 0.30, the composite higher-order LD captures a more constrained baseline of structural dependencies, initiating at a lower absolute magnitude but maintaining structured decay across physical distance. This distinct baseline confirmed that higher-order metrics capture orthogonal topological signals rather than merely mirroring pairwise background linkage. Given that effective double reduction (*α*_*eff*_) introduces localized meiotic noise via sister chromatid co-migration, we hypothesized that regions with elevated DR would spatially antagonize these structured allelic dependencies. By mapping local median *α*_*eff*_ against macroscopic 1-Mb genomic bins, we observed a striking spatial antagonism (Figure 4B). Regions experiencing higher frequencies of effective double reduction demonstrated a substantial, monotonic attenuation in mean higher-order LD intensity, highlighting the disruptive role of DR in modifying the spatial organization of higher-order allelic dependencies.

**Figure 4.**
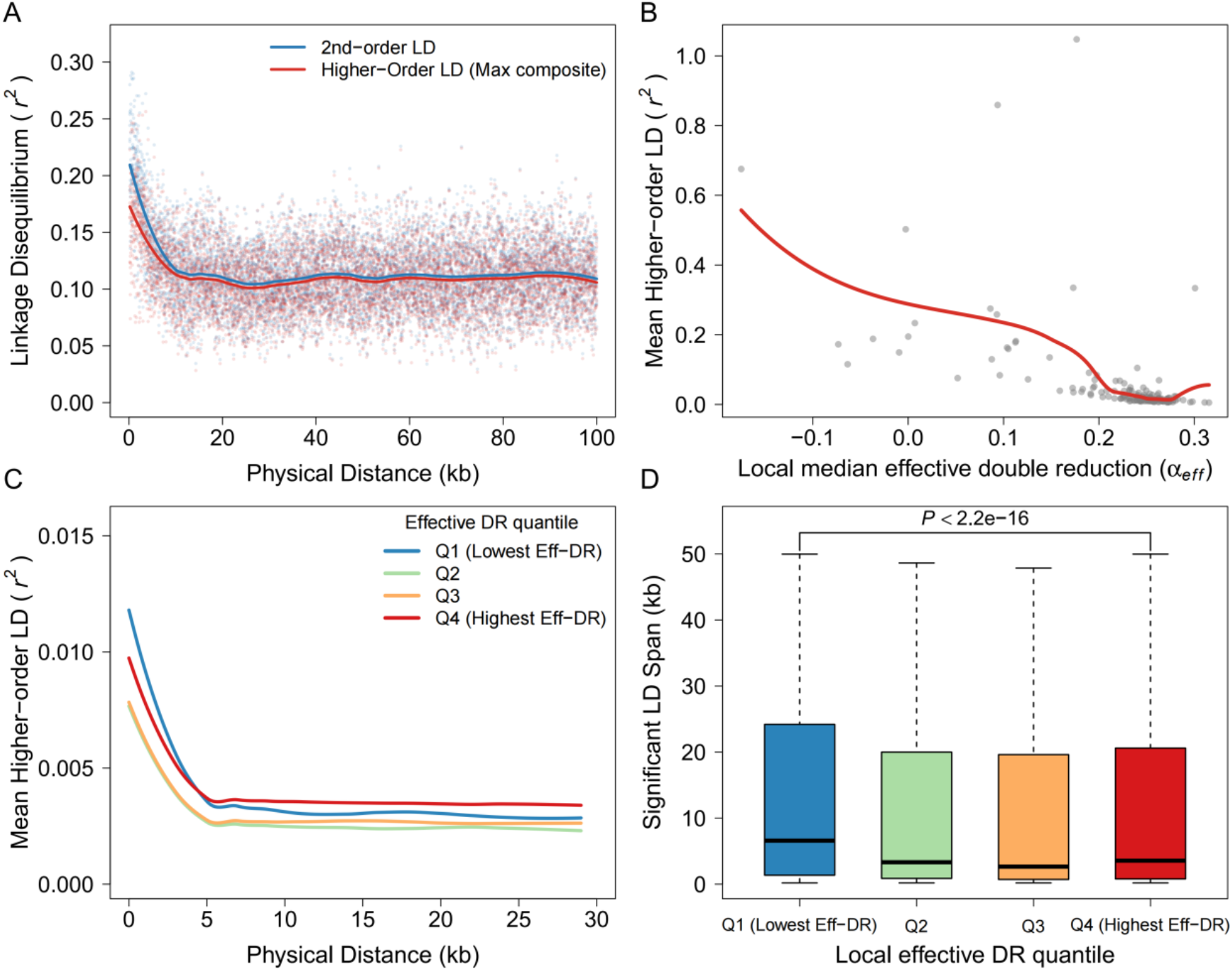
Double reduction is associated with reduced higher-order LD and compressed recombination scales. (A) Genome-wide LD spatial decay in *A. arenosa*. The blue curve represents second-order LD (2nd-order LD), the red curve represents the composite higher-order LD (Higher-Order LD). (B) Relationship between local effective double reduction (*α*_*eff*_) and higher-order LD intensity across 1-Mb windows. (C) Higher-order LD decay stratified by local α_eff_ quantiles (Q1-Q4), where genomic windows are grouped from low (Q1) to high (Q4) effective double reduction. (D) Distribution of significant LD spans across DR regimes.

To rigorously test whether this topological disruption is dose-dependent, we stratified the genome-wide LD decay trajectories by local effective DR quantiles (Figure 4C). The stratified decay analysis provided direct support for our initial hypothesis: areas within the lowest decile of DR (Q1, the blue curve) maintained strong higher-order linkage across considerable physical distances. In contrast, genomic segments that underwent the most pronounced DR (Q4, the red curve) exhibited a swift, nearly instantaneous dissolution of structural LD. This observation underscores how augmented multivalent pairing actively interferes with long-range topological associations.

Consequently, this dose-dependent spatial antagonism dictates the macroscopic boundaries of the effective recombination scale (Figure 4D). We extracted the physical spans of all significant topological edges and compared their distributions across the DR regimes. The physical reach of significant LD was drastically compressed in high-DR regions. While low-Eff-DR environments (Q1) permitted long-range allelic dependencies extending up to ∼50 kb, the maximum physical span in high-Eff-DR environments (Q4) was severely truncated to predominantly under 15 kb. This overwhelming scale compression (Wilcoxon rank-sum test, *P* < 2.2 × 10^-6^) provides definitive quantitative proof that elevated multivalent pairing physically truncates the reach of allelic dependencies. These results demonstrate that meiotic double reduction does not merely alter localized allele frequencies, but acts as a primary architectural sculptor that physically shatters and compresses higher-order allelic networks in autopolyploid genomes.

## Discussion

Traditional polyploid population genetics has long been constrained by the “second-order covariance” framework and the “diploid approximation” paradigm (Meirmans *et al*., 2018; Gerard, 2021). However, the dosage configurations inherent to polyploids drive an exponential expansion of the genotype space, causing fundamentally distinct biological states (e.g., A-A-b-b versus A-a-A-a) to generate nearly indistinguishable statistical signals under conventional pairwise metrics (Blischak *et al*., 2018). Our study advances polyploid population genetics by introducing a tensor-based framework that captures higher-order LD interactions beyond traditional pairwise correlations. This framework enables the accurate modeling of complex multi-allelic dependencies in autopolyploid genomes, providing a robust mathematical pathway for estimating effective double reduction (*α*_*eff*_) and understanding the genetic architecture of polyploid species. We demonstrate that allelic dependencies in polyploid genomes are not merely simple pairwise connections, but rather form a highly interconnected higher-order topological network. This structured dependency implies that specific dosage configurations are co-maintained during population evolution, and their joint distributional deviations cannot be mathematically decomposed into simple combinations of second-order terms (Barton & Turelli, 1991). As evidenced by our genome-wide scans, these higher-order topological metrics successfully capture structured dependency hubs (Kemppainen *et al*., 2015) that are orthogonal to the pairwise background. These long-range hubs not only exhibit elevated network transitivity but also reveal the underlying genomic architecture that remains invisible to traditional second-order LD.

Double reduction (DR) has been narrowly viewed as a cytological byproduct of multivalent meiosis (Lloyd & Bomblies, 2016). Under ideal Hardy-Weinberg equilibrium (HWE), our model provides a perfectly unbiased estimate of pure cytological DR. However, in real natural populations (such as *A. arenosa*), severe Hardy-Weinberg disequilibrium (HWD) statistically confounds the true DR signal with identity-by-descent enrichment, selective sweeps, or non-random mating (Hardy, 2016; Monnahan *et al*., 2019). To encapsulate these compounded frequency distortions, we formalize the global parameter of effective double reduction *α*_*eff*_. A major theoretical correction provided by our study illuminates the hierarchical orthogonal structure of the likelihood space: although severe frequency distortions induced by HWD are systematically absorbed by *α*_*eff*_ (pushing empirical false positive rates toward 100%), the estimation of LD parameters exhibits remarkable asymptotic unbiasedness. Precisely because the allelic association components occupy a subspace orthogonal to the deviations absorbed by *α*_*eff*_, our joint likelihood formulation achieves a rigorous statistical decoupling of pure meiotic mechanisms from complex linkage-driven signals, even under severe population stratification.

Our spatial landscape analysis offers a revolutionary quantitative perspective on the interplay between DR and LD, DR is not merely a nuisance parameter for local allele frequencies; it acts as the primary “physical sculptor” of polyploid genomic topology (Wu et al., 2021; Bomblies, 2020). We quantitatively demonstrate that regions with elevated *α*_*eff*_ highly efficiently shatter long-range LD, drastically compressing the physical span of significant genetic interactions from ∼50 kb in low-Eff-DR regions to predominantly under 15 kb (*P* < 2.2 × 10^-16^). This phenomenon reveals the underlying micro-physical mechanism: complex crossovers and sister chromatid co-migration introduced by multivalent pairing can “dissolve” higher-order topological connections of multiple alleles at an exceptionally high rate (Lloyd & Bomblies, 2016). Consequently, multivalent behavior directly dictates the macroscopic boundaries of the effective recombination scale, sculpting the highly specific recombination hotspot distributions characteristic of autopolyploid genomes.

By contrasting a sexually reproducing natural population (*A. arenosa*) with a clonally propagated cultivated population (*S. tuberosum*), we reveal the absolute dominance of reproductive mode over the topological fate of the genome (McKey *et al*., 2010). Sexual reproduction continuously introduces structural topological perturbations via multivalent meiosis; conversely, sustained clonal propagation bypasses meiosis entirely. Here, we formalize this state as the “Heterozygosity Lock,” an evolutionary phenomenon in which sustained clonal reproduction maintains extreme heterotic advantage by freezing advantageous allelic combinations across generations (Hardigan *et al*., 2017), thereby circumventing the erosive effects of meiotic recombination. This “Lock” causes the potato genome to exhibit a substantial baseline collapse of *α*_*eff*_, with the median plummeting to - 0.71. This reproductive strategy directly precipitates the complete dismantling of higher-order topological networks. Although potato retains limited second-order linkage signals, complex higher-order dependency modules are virtually absent. This finding not only extends the theories of Bourke *et al*. (2015) but also provides quantitative proof that, absent the periodic shuffling of meiotic recombination and segregation, higher-order polysomic networks will inherently disintegrate over clonal evolutionary timescales.

Current mainstream polyploid studies rely heavily on haplotype networks or composite second-order estimators for inference (Gerard, 2021; Zheng *et al*., 2016). However, a fundamental theoretical constraint of these conventional frameworks is the statistical confounding between gametic LD (*D*_*ab*_) and nongametic LD (*D*_*a/b*_). In autopolyploid populations, these two distinct components of allelic association are linearly collapsed within the second-order covariance structure, forcing current estimators to rely on “composite LD” measures that mask the underlying inheritance patterns. Our benchmarking profoundly exposes the vulnerability of this composite approach: as biological complexity (HOLD + DR) and statistical ambiguity (dosage uncertainty) increase (Blischak *et al*., 2018), conventional frameworks exhibit severe systematic overestimation (relative bias >0.25). By leveraging the higher-order moments within our tensor framework, AutoLD provides a more robust mathematical pathway to disentangle these components. However, we acknowledge that the clear separation of *D*_*ab*_ and *D*_*a/b*_ is limited without experimental phasing, even in a more complex setting. This constraint will be overcome as long-read sequencing becomes more common, allowing for the creation of fully phased polyploid reference genomes. This will then enable the full use of our theoretical model’s ability to make inferences.

Understanding the interplay between higher-order allelic topologies and recombination landscapes holds profound strategic implications for modern polyploid genetics and molecular breeding (Bourke *et al*., 2017). In the realm of genome-wide association studies (GWAS), our findings suggest that the chronic “missing heritability” or weak association signals often observed in polyploids are highly likely the consequence of misapplying second-order LD to approximate complex, higher-order polysomic traits. The true causal variants may very well be concealed within highly centralized, higher-order LD hubs (Kemppainen *et al*., 2015). Furthermore, in molecular breeding practices, marker-assisted selection for sexual crops versus clonal crops must adopt fundamentally different marker density and linkage strategies, as their underlying LD topological networks have fundamentally diverged (McKey *et al*., 2010; Gerard *et al*., 2021). Ultimately, the higher-order tensor theory and spatial antagonism principles established in this study provide an indispensable statistical mathematical key for unlocking the complex evolutionary codes of polyploid genomes and accelerating precision crop improvement.

## Methods

### Population-genetic framework and scope

We propose a unified probabilistic framework for jointly inferring double reduction (DR) and two-locus allelic associations in autopolyploid populations. Focusing on biallelic markers, we develop explicit models for autotetraploids (4x) and autohexaploids (6x) to characterize dosage-dependent linkage disequilibrium arising from the segregation of multiple homologous chromosomes under polysomic inheritance. We extend the classical concept of linkage disequilibrium to account for non-random allelic associations within diploid (4x) and triploid (6x) gametes. Although the underlying modeling strategy is shared across ploidy levels, the mathematical formulations for 4x and 6x are derived separately to reflect their distinct meiotic configurations and gametic outcomes. This separation ensures that both the double reduction parameter and higher-order allelic association coefficients retain clear and ploidy-specific biological interpretations.

### Population-genetic assumptions and effective parameters

Our theoretical framework is derived under a classical population-genetic model assuming an infinite, randomly mating population at equilibrium, in which zygotic genotypes arise from the random fusion of gametes. In this idealized situation, zygotic genotype frequencies *z* are expressed as functions of allele frequencies *p* and meiotic segregation parameters, including the double reduction rate *ααα* and allelic association coefficients, rather than treated as free parameters. Under these theoretical conditions, deviations from Hardy-Weinberg equilibrium (HWE) arise exclusively from the cytological occurrence of double reduction inherent to polysomic inheritance.

In natural and cultivated polyploid populations, however, these assumptions are often violated. Processes such as inbreeding, clonal propagation, and population stratification may generate genome-wide Hardy-Weinberg disequilibrium (HWD) independent of meiotic segregation mechanisms. The direct incorporation of all such processes would substantially expand the parameter space and reduce statistical identifiability. To maintain a tractable inference framework, we therefore treat the randomly mating model as a theoretical baseline. When the model is fitted to empirical data, the resulting parameter estimates should be interpreted as effective parameters that account for deviations from the idealized assumptions. We specifically refer to the empirically derived double-reduction coefficient as the effective double reduction rate (*αα*_*eff*_). Under this parameterization, *α*_*eff*_ represents the value of the double-reduction parameter that best explains the observed multilocus genotype frequencies under the theoretical model, while accommodating background deviations from Hardy-Weinberg equilibrium present in real populations. We emphasize that *α*_*eff*_ represents an effective population-genetic parameter inferred from genotype frequency distortions, rather than a direct estimate of cytological double reduction rates.

The framework accommodates varying levels of marker information, including fully informative genotypes with exact dosage calls and partially informative genotypes with ambiguous dosages, such as those arising from low sequencing depth or dominant markers. Observed data *Y Y*are modeled as probabilistic emissions from latent genotypic states *ZG*, allowing inference to proceed even when dosage information is incomplete or conflated.

### Modeling meiotic segregation and double reduction

We model meiotic gamete formation by quantitatively parameterizing DR, a defining feature of polysomic inheritance in which sister chromatids may co-segregate into the same gamete. The extent of DR at a locus is captured by a coefficient *α* ∈ [0, 1], representing the cytological double-reduction rate under the theoretical segregation model.Under this idealized framework, deviations from HWE arise solely from double reduction. When the model is fitted to empirical population data, however, additional sources of HWD may be present due to factors such as inbreeding, clonal propagation, or population structure. As a result, the corresponding parameter estimate is interpreted as an effective parameter (*α*_*eff*_), reflecting both the underlying meiotic segregation process and departures from the idealized assumptions described above.

For clarity, the following derivations are presented under the theoretical segregation model described above. In autotetraploids, meiosis produces diploid gametes. We represent the vector of gametic genotype frequencies, *q*, as a convex combination of two limiting segregation regimes: random chromosome segregation (RCS) and complete equational segregation (CES). Under random mating, the expected gametic distribution is given by

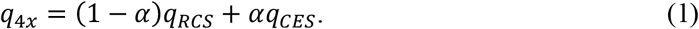

Here,

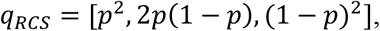

corresponds to binomial sampling under random chromosome segregation, whereas

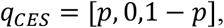

represents the equational segregation limit in which only homozygous gametes are produced as a consequence of complete double reduction. Equivalent closed-form expressions for the resulting gamete frequencies are provided in Supplementary Methods (Section S1.1).

In autohexaploids, meiosis yields triploid gametes, and gametic output depends explicitly on the parental allele dosage. Let *k* ∈ {0, …,6} denote the number of reference alleles in the parental genotype, and let *i* ∈ {0, 1, 2, 3} denote the corresponding dosage in the gamete. The segregation probability *P*(*i*∣*k*) is modeled as a mixture of segregation mechanisms:

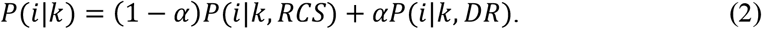

Under RCS, *P*(*i*∣*k*, RCS) follows a hypergeometric distribution corresponding to sampling three chromosomes from six without replacement, reflecting standard polysomic inheritance. The term *P*(*i*∣*k*, DR) captures segregation outcomes associated with double reduction events involving sister chromatid co-segregation. Population-level gametic frequencies are subsequently obtained by marginalizing over the equilibrium distribution of parental genotypes. Explicit expressions are given in Supplementary Methods (Section S1.2)

### Modeling linkage disequilibrium through allelic association structures

We define linkage disequilibrium (LD) as the deviation of the joint gametic frequency from the product of the corresponding marginal frequencies. In contrast to diploid systems, where LD is fully characterized by a single parameter, gametes in autopolyploids carry multiple alleles per locus, giving rise to a structured set of dosage-dependent association terms.

In autotetraploids, gametes are diploid. Let *P*(*A*_*i*_*B*_*j*_) denote the joint probability that a gamete carries dosage *i* at locus *A* and dosage *j* at locus *B*. We express this joint distribution as

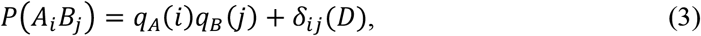

where *q*_*A*_(*i*) and *q*_*B*_(*j*) are the marginal gametic frequencies derived under the single-locus segregation model, and *δ*_*ij*_(*D*) aggregates contributions from digenic, trigenic and quadrigenic associations arising from the co-segregation of homologous chromatids. The deviation term is parameterized by a set of LD coefficients *D*, which capture allelic associations at increasing orders of interaction, ranging from pairwise associations between single alleles at the two loci to joint associations involving the full diploid gametic states. This decomposition provides a complete and non-redundant representation of dosage-dependent LD in autotetraploid gametes.

In autohexaploids, gametes are triploid, substantially expanding the space of possible allelic associations. For *i, j* ∈ {0, 1, 2, 3}, the joint distribution *P*(*A*_*i*_*B*_*j*_) is modeled using an analogous decomposition, with the deviation term *δ*_*ij*_(*D*) incorporating higher-order interaction components that arise from the joint segregation of three alleles per locus. These components include deviations in triploid gamete composition from random sampling expectations, as well as between-locus interaction terms that couple allele dosage states across loci, up to the level of the full triploid configuration at both loci. This parameterization generalizes classical diploid LD by accommodating the complex, dosage-dependent association structure induced by polysomic inheritance. The analytically derived of linkage disequilibrium for autotetraploids and autohexaploids are provided in Sections 2.1 and 2.2 of the Supplementary Methods, respectively.

### Parameter estimation from fully and partially informative genotypes

Model parameters are estimated from observed two-locus genotype counts under either fully informative or partially informative marker data. Under fully informative markers, observed genotypes map uniquely to dosage-defined zygotic classes (25 for autotetraploids and 49 for autohexaploids). In this setting, estimation proceeds by directly reconstructing the two-locus gametic frequency distribution under the assumption of random mating. These gametic frequencies are then mapped to allele frequencies, double-reduction coefficients, and linkage disequilibrium components using the closed-form population-genetic transformations derived in Supplementary Methods Sections 2.1 and 2.2.

For partially informative markers (e.g., due to low sequencing depth or dominant encoding), multiple latent zygotic states are conflated into a single observed phenotype. Let *Z* denote the latent two-locus zygotic genotype and *Y* the observed marker category. The probability of observing an observed genotype *y* is modeled as a probabilistic mixture:

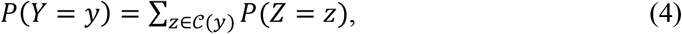

where 𝒞 (*y*) represents the set of latent zygotic configurations compatible with observation. we employ an Expectation-Maximization (EM) algorithm to reconstruct the underlying two-locus gametic frequency distribution under the assumption of random mating. The EM procedure is used solely to recover latent frequency tables rather than population-genetic parameters directly.

Allele frequencies and linkage disequilibrium coefficients are subsequently obtained from the reconstructed frequency tables through analytical transformations, and the double reduction coefficient (*α*) is then determined by solving the marginal constraints implied by the reconstructed table. When applied to empirical population data, this estimate is interpreted as the effective double reduction parameter (*α*_*eff*_ *α*), reflecting both the underlying meiotic segregation process and potential deviations from Hardy–Weinberg equilibrium in real populations.This separation ensures that inference is anchored in biologically interpretable population-genetic quantities, rather than in algorithm-specific optimization parameters. The detailed EM formulations and parameter mapping functions for both autotetraploids and autohexaploids are provided in Supplementary Methods Section 3.

### Hypothesis testing for parameter significance

To assess the statistical significance of individual population-genetic parameters, we employed a likelihood-ratio test (LRT) framework. Following the parameter estimation described above, let 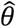 denote the maximum likelihood estimate of the full parameter vector obtained via the EM algorithm. This estimate corresponds to the global maximum of the likelihood function, 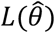. For a specific parameter of interest 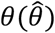, we formulated the hypothesis test as:

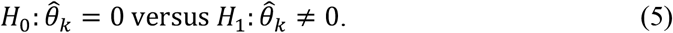

To evaluate *H*_0_, we calculated the restricted maximum likelihood estimate, 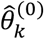, by maximizing the likelihood function subject to the constraint *θ*_*k*_ = 0 while profiling out all remaining nuisance parameters. The test statistic is defined as:

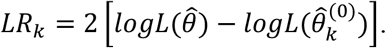

This statistic quantifies the reduction in model fit when the parameter *θ*_*k*_ is removed. Under the null hypothesis and standard regularity conditions, *LR*_*k*_ follows an asymptotic chi-square distribution with one degree of freedom.

### Normalization of disequilibrium coefficients

The allelic association parameters estimated by our model are covariance-based measures whose magnitudes depend on allele frequencies. Raw values of disequilibrium coefficients are not directly comparable across loci or populations with different allele frequency spectra. To facilitate comparison, we extend Lewontin’s normalization (Lewontin, 1964) for gametic linkage disequilibrium to the hierarchy of dosage-dependent association components considered here.

Let *D* denote a generic disequilibrium component, including digenic, trigenic, or higher-order association terms. Conditional on the observed allele frequencies (*p*A, *p*B), the magnitude of *D* is constrained by the requirement that all implied two-locus gametic frequencies remain within the probability simplex. We, yielding the normalized disequilibrium measure

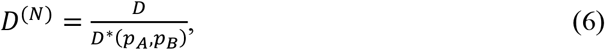

where the scaling constant *D*^*^(*p*_A_, *p*_B_) represents the maximal attainable magnitude of *D* given its sign. Following Lewontin’s *D*^<^ normalization, this bound is approximated using marginal-frequency-based limits:

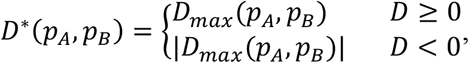

where the limiting values are derived from feasible combinations of marginal gametic frequencies.

For higher-order association components in polyploid systems, the limiting bounds are approximated using marginal frequency proxies corresponding to the effective dosage order of the association term. This approximation provides a stable scaling factor while avoiding the computational complexity of solving the full probability-simplex constraints. The resulting normalized coefficient *D*^(*N*)^ lies in the interval [-1, 1] and is used for visualization and cross-locus comparison of disequilibrium magnitudes.

### Comparative evaluation of pairwise LD estimation methods

To evaluate the performance of the proposed joint inference framework (AutoLD) for estimating pairwise linkage disequilibrium (*D*_*eab*_), we compared it with three commonly used approaches representing different levels of model simplification.The diploid approximation (Dip) collapses autopolyploid dosage genotypes into pseudo-diploid states, ignoring polysomic inheritance and allele dosage variation. Pairwise LD is then calculated using standard diploid formulations.The haplotype EM approach (Hap) estimates two-locus haplotype frequencies using a standard expectation-maximization algorithm and computes LD as *D*_*eab*_ = *P*_*AB*_ - *p*_*A*_*p*_*B*_. This formulation inherently assumes random chromosome pairing (α = 0) and the absence of higher-order gametic associations.We further included the composite LD estimator implemented in the R package ldsep (Gerard, 2021), which infers pairwise LD directly from genotype data. While it accommodates dosage uncertainty, it does not explicitly parameterize higher-order LD or double reduction.Because these three approaches do not model higher-order gametic associations or segregation distortions, the resulting estimates essentially correspond to a composite pairwise LD.

### Empirical data used for model evaluation

To evaluate the proposed framework under realistic population-genetic conditions, we analyzed two autotetraploid populations with dosage-resolved SNP genotypes. These datasets represent distinct biological contexts, including a natural population and a cultivated crop population, while sharing the key property that allele dosage can be inferred with high confidence, enabling direct application of the proposed polyploid linkage disequilibrium framework.

#### *Arabidopsis arenosa* population

The first dataset was derived from a natural population of *Arabidopsis arenosa*, a well-established autotetraploid model species for studying polysomic inheritance and genome evolution in polyploids (NCBI SRA accession PRJNA484107; Monnahan *et al*., 2019). Raw reads were aligned to the reference genome using BWA (Li, 2013) and duplicate reads were removed using Picard tools (Picard Toolkit, 2019). Variants were identified with GATK HaplotypeCaller (McKenna *et al*., 2010), specifically adapted for polyploid genotyping by incorporating allele dosage estimation. Variant filtering was applied to retain high-confidence biallelic SNPs. Specifically, variants were required to satisfy the following criteria: minimum mapping quality ≥ 30, minimum base quality ≥ 20, and minimum read depth ≥ 5 per individual. Sites with a missing genotype rate greater than 10% or a minor allele frequency (MAF) below 0.01 were excluded. Allele dosage was inferred from genotype likelihoods produced by the variant-calling pipeline, and only SNPs with confidently resolved allele dosage were retained for downstream analyses. After filtering, 170 individuals and 170,806 SNPs remained for analysis.

#### Cultivated potato population

The second dataset consisted of a cultivated autotetraploid potato (*Solanum tuberosum*) population (NGDC accession GVM000314). In this dataset, genotype information was provided as VCF files containing genotype probabilities derived from polyploid genotyping. Allele dosage was inferred from these genotype probabilities by selecting the most likely genotype configuration for each individual. Quality-control filtering was applied to remove low-confidence variants and sites with excessive missing data. Filtering criteria included a maximum missing rate of 10% and a minor allele frequency threshold of 0.01. After filtering, 527 individuals and 4,333,977 SNPs were retained for analysis.

### SNP thinning and non-parametric modeling of LD decay

To mitigate the confounding effects of extreme local linkage and reduce computational redundancy in genome-wide LD estimation, we implemented a physical distance-based SNP thinning strategy. SNPs within each chromosome were ordered by their genomic coordinates, and a sequential filtering algorithm was applied to enforce a minimum physical distance threshold between adjacent retained SNPs. This thinning procedure resulted in a more uniformly distributed subset of markers, minimizing localized autocorrelation while preserving the global genomic architecture for subsequent higher-order two-locus association analyses.

Genome-wide linkage disequilibrium decay was evaluated by examining the relationship between physical distance and allelic association strength among marker pairs. Pairwise LD statistics were computed for SNP pairs located on the same chromosome. In addition to classical second-order LD, higher-order two-locus association components were also calculated under the proposed framework.To characterize the distance-dependent decay patterns, marker pairs were grouped into discrete physical distance bins of 1000 bp. For each bin, the mean *r*^2^ value was computed to summarize LD magnitude. Continuous LD decay trajectories were then modeled using a non-parametric locally estimated scatterplot smoothing (LOESS) regression applied to the binned mean *r*^2^ values (span = 0.25), allowing flexible capture of non-linear decay dynamics without imposing parametric assumptions. Decay profiles were visualized under two complementary analytical resolutions. A granular representation independently traces the decay trajectories of individual higher-order LD components, whereas a composite representation contrasts the classical second-order LD against the maximal magnitude of all higher-order association terms. To focus on the biologically relevant scale of recombination-driven LD breakdown, the genome-wide decay curves were truncated at a maximum physical distance threshold.

### Statistical analysis of spatial clustering and topological LD

To rigorously validate the spatial clustering of higher-order LD, we implemented a matched-window permutation framework across non-overlapping 100-kb genomic bins. The observed edge densities were evaluated against an empirical null distribution generated via chromosome-wise shuffling of binary significance labels (*N* = 100-200). This procedure disrupts local spatial architecture while preserving global interaction prevalence, ensuring that the observed clustering reflects biological signals rather than random variation. Spatial clustering intensity was quantified using a standardized *Z*-score:

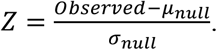

Hotspots were defined conservatively at the extreme upper tail (*Z* ≥ 5), after excluding low-information windows (< 100 tested SNP pairs) to prevent variance inflation. This step ensured that only robust spatial clustering signals were considered, providing a reliable measure of LD hotspot intensity. Finally, to assess the topological antagonism driven by multivalent pairing, LD pairs were stratified into quantiles based on local median effective double reduction (*α*_*eff*_) values. The physical spans of significant topological edges across these DR regimes were statistically compared using the non-parametric Wilcoxon rank-sum test, ensuring that any observed differences reflected biologically meaningful changes in the LD landscape rather than random variation.

## Data and code availability

The empirical datasets analyzed during the current study are publicly available. The sequencing data for the natural population of the autotetraploid model species *Arabidopsis arenosa* can be accessed in the NCBI Sequence Read Archive (SRA) under the accession number PRJNA484107. The data for the cultivated autotetraploid potato (*Solanum tuberosum*) population are available in the National Genomics Data Center (NGDC) under the accession number GVM000314. The AutoLD R package developed in this study, along with its source code, detailed tutorials, and example datasets, is open-source and freely available on GitHub at https://github.com/SDUT-Sysbio/AutoLD.

## Acknowledgements

This work was supported by the National Natural Science Foundation of China (Grant Nos. 32070601 to L.J. and 32370689 to F.Y.).

## Author contributions

L.J. and F.Y. conceived and supervised the project. W.D. derived the mathematical models, performed the computer simulations, and conducted the real data analysis.Z.L. and D.J. assisted with the empirical data analysis. C.Z. carried out the bioinformatics analyses. W.D., N.W., and L.J. wrote the original draft of the manuscript. L.J. and F.Y. critically reviewed and edited the manuscript. All authors read and approved the final manuscript.

